# Sex-based disparities in DNA methylation and gene expression in late-gestation mouse placentas

**DOI:** 10.1101/2023.09.23.559106

**Authors:** Lisa-Marie Legault, Mélanie Breton-Larrivée, Alexandra Langford-Avelar, Anthony Lemieux, Serge McGraw

**Affiliations:** CHU Ste-Justine Research Center, 3175 Chemin de la Côte-Sainte-Catherine, Montréal, QC H3T 1C5, Canada; Department of Biochemistry and Molecular Medicine, Université de Montréal, 2900 Boulevard Edouard-Montpetit, Montréal, QC H3T 1J4, Canada; Department of Obstetrics and Gynecology, Université de Montréal, 2900 Boulevard Edouard-Montpetit, Montréal, QC H3T 1J4, Canada

**Keywords:** Placenta, DNA methylation, gene expression, late pregnancy, mouse, sex differences, development

## Abstract

**Background:** The placenta is vital for fetal development and its contributions to various developmental issues, such as pregnancy complications, fetal growth restriction, and maternal exposure, have been extensively studied in mice. The placenta forms mainly from fetal tissue and therefore has the same biological sex as the fetus it supports. Extensive research has delved into the placenta’s involvement in pregnancy complications and future offspring development, with a notable emphasis on exploring sex-specific disparities. However, despite these investigations, sex-based disparities in epigenetic (*e.g*., DNA methylation) and transcriptomic features of the late-gestation mouse placenta remain largely unknown.

**Methods:** We collected male and female mouse placentas at late gestation (E18.5, *n* = 3/sex) and performed next-generation sequencing to identify genome-wide sex differences in transcription and DNA methylation.

**Results:** Our comparison between male and female revealed 358 differentially expressed genes (DEGs) on autosomes, which were associated with signaling pathways involved in transmembrane transport and the responses to viruses and external stimuli. X chromosome DEGs (*n* = 39) were associated with different pathways, including those regulating chromatin modification and small GTPase-mediated signal transduction. Differentially methylated regions (DMRs) were more common on the X chromosomes (*n* = 3756) than on autosomes (*n* = 1705). Interestingly, while most X chromosome DMRs had higher DNA methylation levels in female placentas and tended to be included in CpG dinucleotide-rich regions, 73% of autosomal DMRs had higher methylation levels in male placentas and were distant from CpG-rich regions. Several DEGs were correlated with DMRs. A subset of the DMRs present in late-stage placentas were already established in mid-gestation (E10.5) placentas (*n* = 348 DMRs on X chromosome and 19 DMRs on autosomes), while others were acquired later in placental development.

**Conclusion:** Our study provides comprehensive lists of DEGs and DMRs between male and female that collectively cause profound differences in the DNA methylation and gene expression profiles of late-gestation mouse placentas. Our results demonstrate the importance of incorporating sex-specific analyses into epigenetic and transcription studies to enhance the accuracy and comprehensiveness of their conclusions and help address the significant knowledge gap regarding how sex differences influence placental function.

**Highlights:**

- In the mouse placenta, sex-specific gene expression and DNA methylation profiles, enriched in various metabolic and developmental pathways, are observed for both X-linked and autosomal genes from mid-gestation onward.
- Regions with different DNA methylation are commonly found in CpG-rich areas on the X chromosomes and in CpG-poor regions on autosomes.
- A subset of the DMRs observed in late-stage placentas were already established in mid-gestation placentas, whereas others were acquired during the later stages of placental development.
- Several DNA methylation sex differences could be correlated with sex differences in gene expression.
- The results highlight the importance of including sex-based analyses in epigenetic and transcriptional studies of the mouse placenta.

**Plain English summary:** The placenta is a crucial organ for a healthy pregnancy and proper fetal development, and its functions are often studied in mice. The placenta stems from the developing embryo, and therefore shares its sex. Male fetuses have higher risks of pregnancy complications and neurodevelopmental disorders, and these risks are linked to placenta functions. However, how the placenta’s sex influences the proteins it contains—and therefore, how it helps the fetus develop—remains largely unknown. We used cutting-edge techniques to systematically examine late-pregnancy mouse placentas, cataloging the genes being expressed (*i.e*., sections of DNA used to make proteins) and the patterns of a specific DNA mark (called methylation) that controls gene expression. We identified several genes with important placental functions, such as protecting the fetus from viruses and responding to environmental changes, whose expression levels were sex-specific. We also observed differences in DNA methylation between male and female placentas. Most DNA methylation differences were on the X-chromosomes associated with sex, and the majority had higher methylation levels in female placentas. Conversely, on other chromosomes, most differences present an increased level of DNA methylation in male placentas. As methylation affects gene expression, we found links between the changes. Additionally, we found that some sex differences in the placenta were already present earlier in pregnancy. Our findings provide important insights into the molecular differences between male and female mouse placentas during late pregnancy. Including sex-specific analyses in placenta studies will improve our understanding of how the placenta ensures the healthy development of male and female fetuses.

## BACKGROUND

Biological sex significantly impacts various aspects of life, ranging from cellular processes to the overall functioning of the organism. The placenta plays a critical role in allocating maternal-fetal resources, including oxygenation, nutrition, and metabolic exchanges between the mother and the fetus. It also acts as a protective barrier, responding to infection, stress, and other external factors to safeguard the developing fetus (1–3). The placenta comprises mainly tissue derived from early embryonic development and shares the same sex chromosomes as the embryo. Still, many studies do not consider the roles of the placenta’s biological sex in their analyses.

Given its importance for fetal development, placental dysfunction and altered responses to external stressors can lead to numerous pregnancy complications, including preeclampsia, fetal growth restriction, gestational diabetes, and preterm birth (4, 5). Recent studies demonstrate that the placenta has sex-specific responses to certain stimuli or perturbations, which influence their impacts on the embryo (6–8). These findings align with existing evidence—primarily based on meta-analyses of human pregnancies— consistently indicating that male fetuses have higher susceptibility to pregnancy complications such as gestational diabetes, premature membrane rupture, preterm birth, and macrosomia (8–14). Males also exhibit a higher prevalence of neurodevelopmental disorders, including dyslexia and attention deficit hyperactivity disorder. The increased occurrence of neurodevelopmental disorders in males has been associated with placental dysfunction and adaptations to diverse pregnancy conditions (6). Nevertheless, substantial evidence directly linking these disorders to placental malfunction remains elusive. Collectively, the evidence indicates that biological sex influences how the placenta functions and adapts to diverse pregnancy conditions; however, how sex-specific epigenetic and transcriptomic responses contribute to these increased risks to male fetuses remains unknown.

The critical roles of DNA methylation and gene expression in regulating development have been studied in detail in various biological systems (15–17). Significant differences in the methylation patterns and gene expression levels of an embryo and its placenta are established quickly after implantation (18, 19). Mouse embryos display sex-specific DNA methylation patterns, which can be differently altered following adverse maternal exposure (*e.g.* alcohol, environmental toxicant, drug) (20–23). While initially thought to occur primarily on sex chromosomes, sex differences in DNA methylation and gene expression have been detected throughout the genome in various tissues and organs, including the human placenta (24–29). Systematically investigating the DNA methylation and gene expression profiles of male and female mouse placentas will provide valuable insights into these sex-based variations in placental functions.

In this study, we have systematically identified DNA methylation and gene expression sex differences between male and female late-gestation mouse placentas. We uncovered numerous disparities in genes linked to important placental functions and embryonic development, including some on autosomal chromosomes. Our findings underscore the importance of including male and female samples and analyzing them independently, as biological sex results in important molecular differences that could profoundly impact a study’s results.

## METHODS

### Mouse studies and tissue collection

All animal studies were approved by the CHU Ste-Justine Research Center *Comité Institutionnel de Bonnes Pratiques Animales en Recherche* under the guidance of the Canadian Council on Animal Care. Male and female C57BL/6 mice (Charles River Laboratories, Wilmington, MA, USA) were housed in a 12 h light/dark cycle with unlimited access to food and water and were mated at 8 weeks old. Females who had developed copulatory plugs by the next morning were considered pregnant with day 0.5 embryos (E0.5) and were separated from the males and housed together. The pregnant mice were euthanized at E10.5 or E18.5, and the placentas were dissected to remove maternal tissue (myometrium and decidua) under Leica Stereo Microscope, flash frozen in liquid nitrogen, and stored at −80°C until DNA and RNA extraction. The sex of each placenta was determined by *Ddx3* qPCR using digested DNA from the corresponding embryo’s tail (21).

### DNA/RNA extraction and library preparation

We selected healthy looking placentas from three different litters (*n* = 3/sex) for DNA and RNA extraction. Whole placentas were homogenized to powder in liquid nitrogen. Samples were split in two, and the halves were used to extract genomic DNA with a QIAamp DNA Micro Kit (Qiagen, Hilden, Germany, #56304) and RNA using a RNeasy Mini Kit (Qiagen #74004), respectively, following the manufacturer’s recommendations. Extracted DNA and RNA were quantified using a Qubit dsDNA BR (Broad Range) Assay Kit (Thermo Fisher Scientific, Waltham, MA, USA, #Q32853) and a High Sensitivity RNA assay kit (Thermo Fisher Scientific, #Q32852), respectively, on a Qubit 3.0 Fluorometer (Thermo Fisher Scientific #Q33217).

Mouse methyl capture sequencing (Methyl-Seq) libraries were generated using the SureSelect*^XT^* Methyl-Seq Target Enrichment System (Agilent, Santa Clara, CA, USA, #G9651B) and the SureSelect*^XT^* Mouse Methyl-Seq target enrichment panel (Agilent, #5191-6704) following the manufacturer’s recommendations. Briefly, 1 µg of genomic DNA was used for library preparation. Target regions were enriched by biotinylated precipitation, followed by sodium bisulfite conversion and library amplification/indexing. Libraries were quantified using a Qubit dsDNA HS (High Sensitivity) Assay Kit (Thermo Fisher Scientific #Q32854) on a Qubit 3.0 Fluorometer. Library quality control was assessed using a BioAnalyzer (Agilent) followed by paired-end sequencing on a NovaSeq 6000 S4 sequencer at the Genome Québec core facility. We obtained 112–139M reads for the sequenced libraries.

Reduced Representation Bisulfite Sequencing libraries were performed based on the rapid RRBS protocol (rRRBS)(19, 21, 31–33). Briefly, 500 ng of DNA was digested with *Msp1* restriction enzyme and adapters were attached to DNA fragments. DNA was converted using sodium bisulfite treatment and amplification/indexation of libraries was performed. Libraries were quantified using QuBit fluorimeter apparel with the High Sensitivity DNA assay kit (ThermoFisher Scientific #Q32854). Quality of the libraries was assessed using BioAnalyzer followed by paired-end sequencing was done on Illumina HiSeq 2500 at the Genome Québec core facility.

We generated mRNA-sequencing (mRNA-Seq) libraries using 500 ng of good-quality RNA (RIN ˃7and a NEBNext Ultra II Directional RNA Library Prep Kit (New England BioLabs, Ipswich, MA, USA, #E7760L) at the Genome Québec core facility and paired-end sequenced on a NovaSeq6000 S4 sequencer. We obtained 26–40M reads for the sequenced libraries.

### Bioinformatics analyses

Post-sequencing bioinformatic analyses of mRNA-Seq data were performed using the GenPipes RNA-Seq pipeline (v4.1.2) (34) with alignment to the mouse GRCm38 genome (mm10). Differential gene expression analysis was performed with the R (v3.5.0) package DESeq2 (v1.24.0) (35), including multiple testing correction using the Benjamini-Hochberg false discovery rate procedure, at a significance threshold of *p* ˂ 0.05 with a normalized read count ≥ 1 in all replicates to get a broad overview of placenta transcriptome, including genes presenting a low level of expression.

Methyl-Seq data were analyzed using the GenPipes Methyl-Seq pipeline (v3.3.0) (34) with reads aligned to the mouse GRCm38 reference genome, and methylation counts were obtained using Bismark (v0.18.1) (36). rRRBS data were analyzed using our custom pipeline (32), including tools such as Trim Galore (v0.3.3) (37), BSMAP (v2.90) (38) and R, with alignment to the mouse GRCm38 reference genome. DMRs for both Methyl-Seq and rRRBS datasets were identified with the R package MethylKit (version 1.8.1) (39) using the Benjamini-Hochberg false discovery rate procedure. Fixed parameters were used, including 100-bp stepwise tiling windows and a threshold of *q* < 0.01. Reported DNA methylation levels represent the average methylation levels of all CpG dinucleotides (CpGs) within a tile for all samples within a condition. The number of CpGs and bisulfite conversion rate (> 96%) of each tile were obtained using a custom Perl script (21).

Genome annotation of the tiles was performed using Homer (version 4.10.1) (40) and the mouse mm10 reference genome. Intragenic regions were defined as all annotations not in promoter or intergenic regions, such as 3’ UTR (untranslated regions), 5’ UTR, exons, introns, TTS (transcriptional termination site), non-coding regions. Gene ontology term enrichment analyses of differentially methylated tiles located in intragenic regions were performed in Metascape (metascape.org) (41). Repeats and CpG island coordinates for the mm10 genome were obtained from the UCSC Genome Browser (genome.ucsc.edu) (42). CpG shores and CpG shelves represent the regions within 0–2 kb and 2–4 kb of CpG islands, respectively, as previously described (19, 21). Statistical analyses were performed in R (v3.5.0) or GraphPad Prism (version 9.5.0; GraphPad Software, San Diego, CA, USA).

## RESULTS

### Late-gestation mouse placentas have sex-specific gene expression profiles

To reveal the molecular differences between male and female placentas during mouse development, we first conducted an in-depth gene expression analysis of six whole E18.5 placentas (*n* = 3/sex, from three different litters) using RNA-Seq. The expressed genes were divided into three groups based on their location on autosomal chromosomes, the X chromosomes, or the Y chromosome. We found that male placentas contained more transcribed autosomal genes than their female counterparts (i.e number of genes expressed with a normalized read counts ˃1 in all 3 replicated of one sex; 26182 vs. 25758; Figure 1A). Female placentas had slightly more transcribed X chromosome genes than male placentas (1139 and 1129 in male and female placentas, respectively; Figure 1A). As expected, Y chromosome gene expression was observed exclusively in male placentas, with seven transcripts detected (Figure S1).

**Figure 1.**
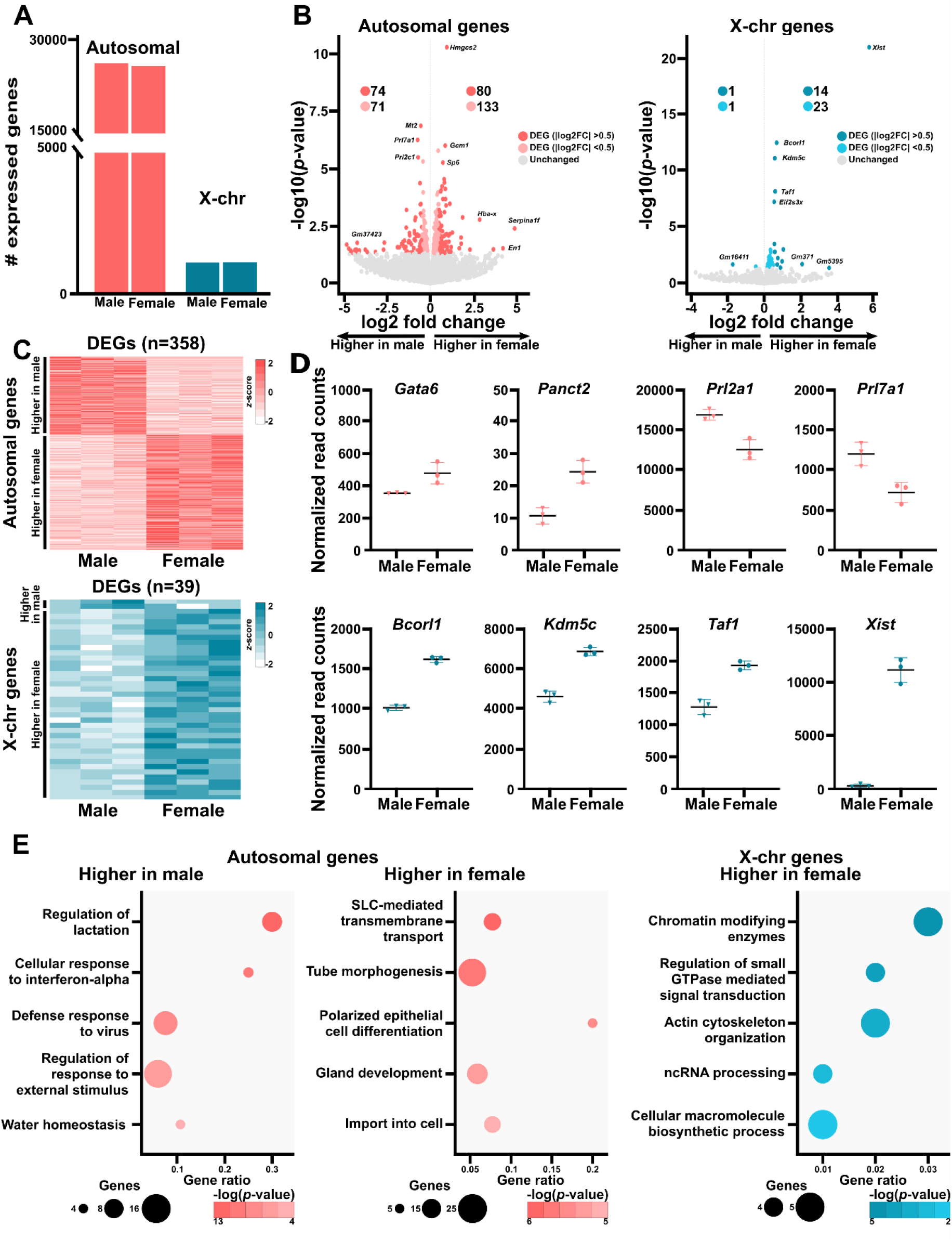
DEGs occur throughout the genomes of late-gestation mouse placentas. A) Genes expressed in male and female E18.5 placentas from the autosomes (male *n* = 26182, female *n* = 25758) and X chromosomes (male *n* = 1126, female *n* = 1139). Genes included had read counts ˃ 0 in all samples of the relevant sex. **B)** Differential expression analysis of autosomal (*n* = 28194; left) and X chromosomal (*n* = 1286; right) genes in male and female placentas. Statistically significant DEGs are represented by colored dots (*p* < 0.05; *n* = 358 autosomal, *n* = 39 X chromosomal). Darker dots indicate genes with |log_2_ fold change| values > 0.5 (*n* = 154 autosomal, *n* = 15 X chromosomal). **C)** Expression levels (*z*-scores) of 358 autosomal (top) and 39 X chromosomal (bottom) DEGs. **D)** Normalized read counts of genes with sex-specific expression. **E)** The top five pathways associated with the autosomal (left and middle) and X chromosome (right) DEGs.

We identified 397 significantly differentially expressed genes (DEGs; *p* < 0.05) between male and female placentas. The majority (358) were located on autosomal chromosomes; of these, 145 were more highly expressed in male placentas and 213 were higher in female placentas (Figure 1B and C; Table 1). The X chromosomes contained 39 DEGs, with two and 37 upregulated in male and female placentas, respectively (Figure 1B and C; Table 2). Strikingly, many of the DEGs on autosomal chromosomes (43%) and X chromosomes (38%) had |log_2_ fold change| values ≥ 0.5, indicating substantial differences in gene expression between male and female placentas from all shared chromosomes (Figure 1B). Most genes (99%) had < 10000 normalized read counts, and approximately 62% (*n* = 18277) had < 100 normalized read counts (Figure S2A–B). DEGs on autosomal or X chromosomes were not restricted to genes with lower normalized read counts but were distributed across the abundance spectrum, as observed for genes without sex-specific expression (Figure S2A–B). DEGs included *Gata6*, *Panct2*, *Prl2a1*, and *Prl7a1* on the autosomes and *Bcorl1*, *Kdm5c*, *Taf1,* and *Xist* on the X chromosomes (Figure 1D).

**Table 1.**
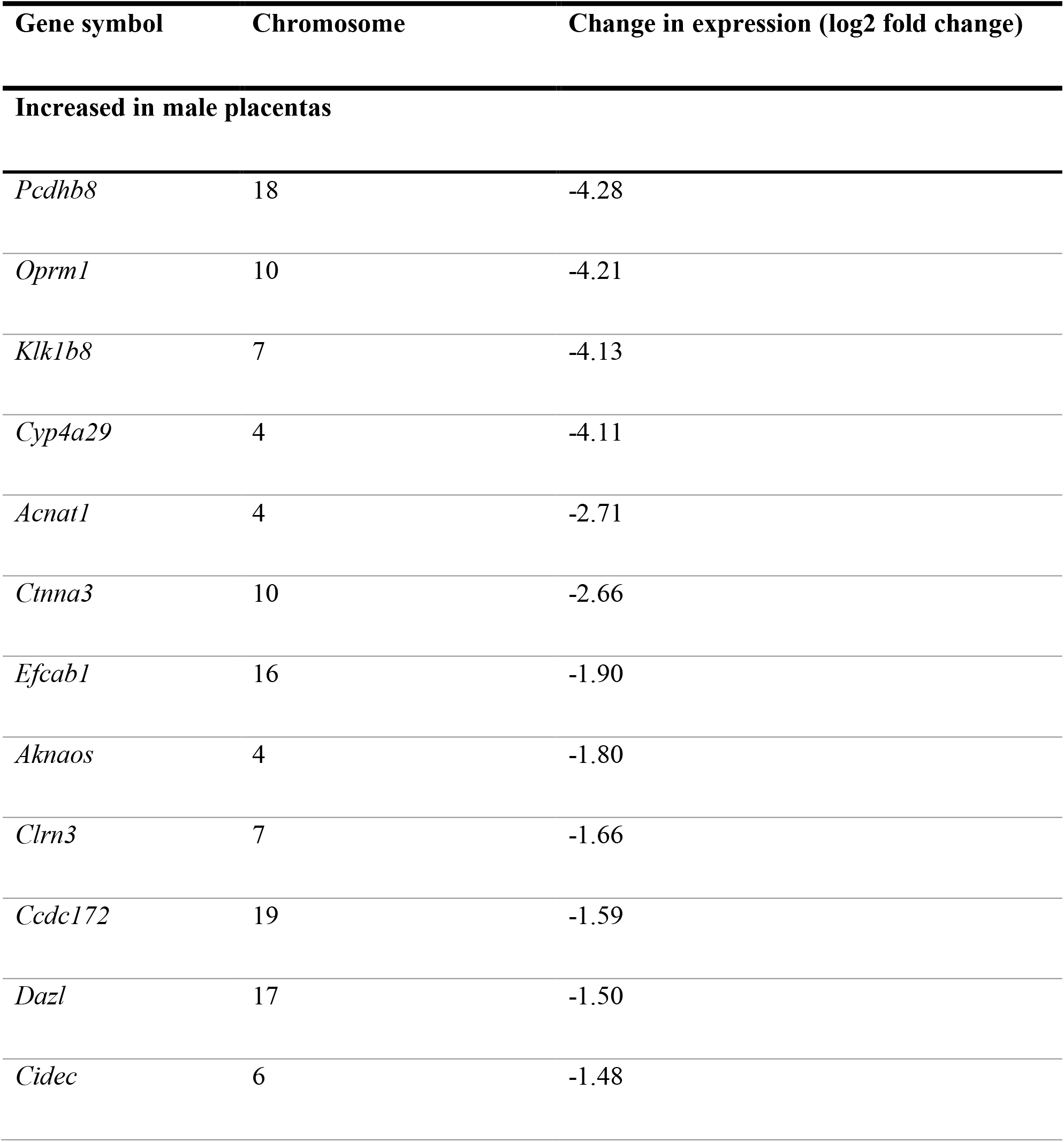
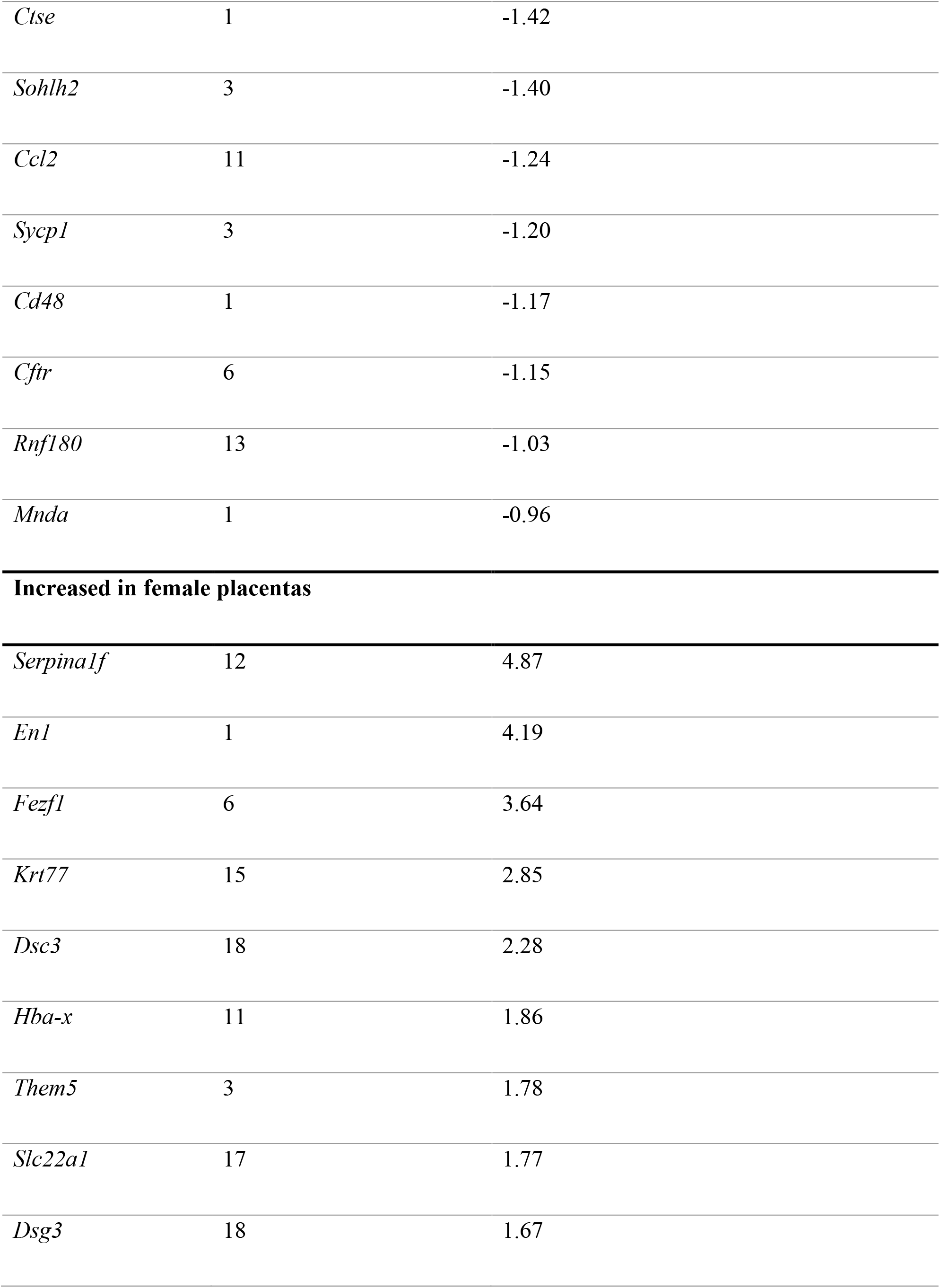
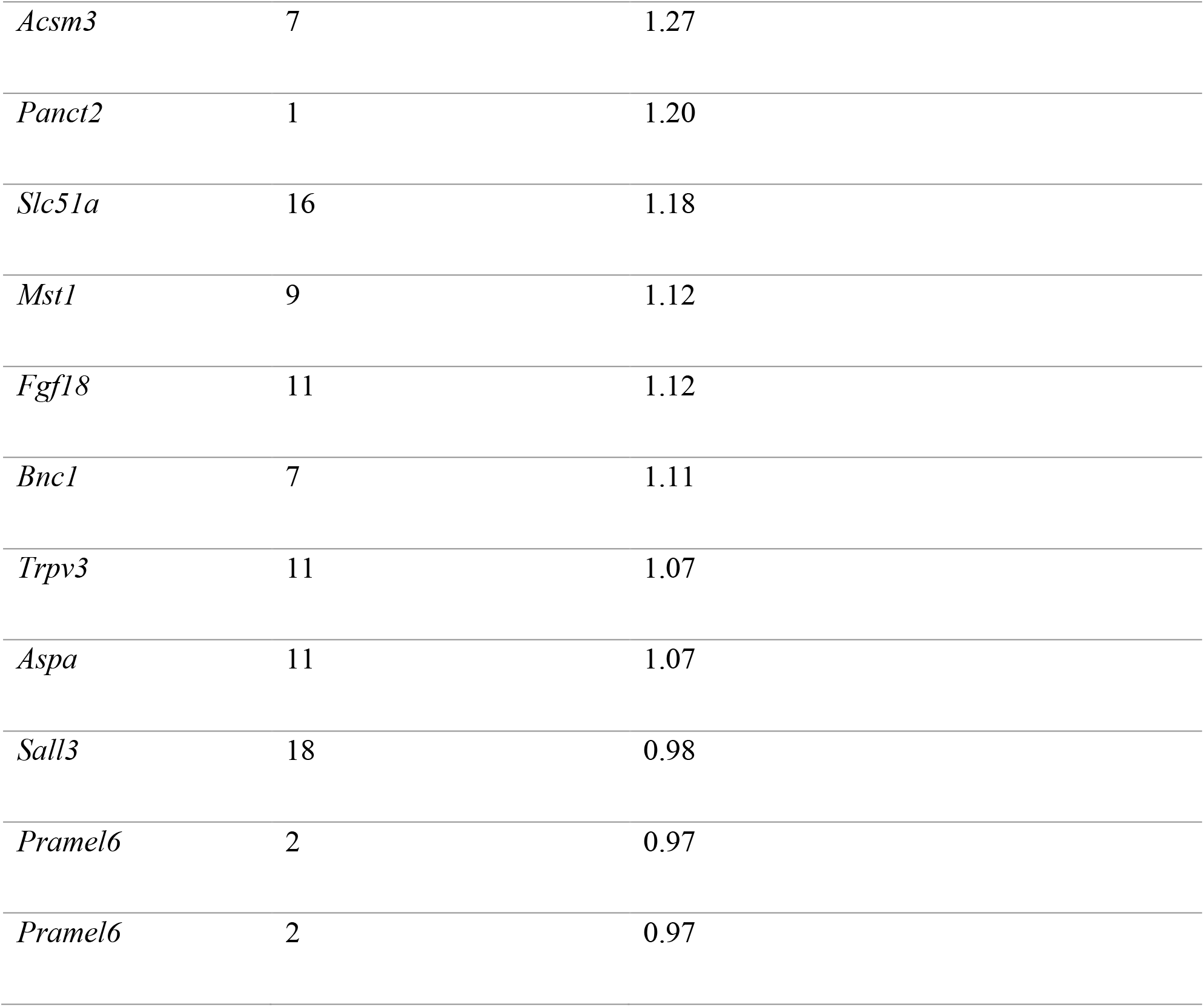
Top DEGs on autosomal chromosomes (*p* ˂ 0.05)

**Table 2.**
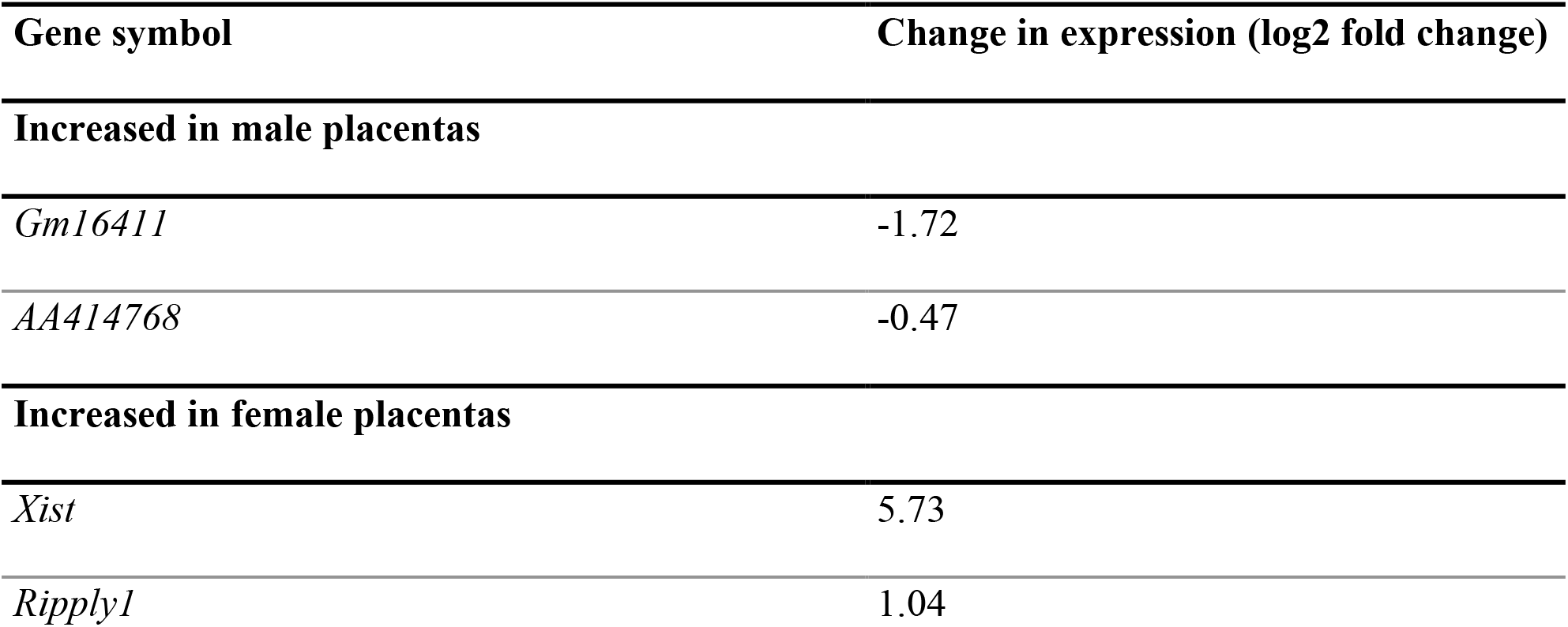
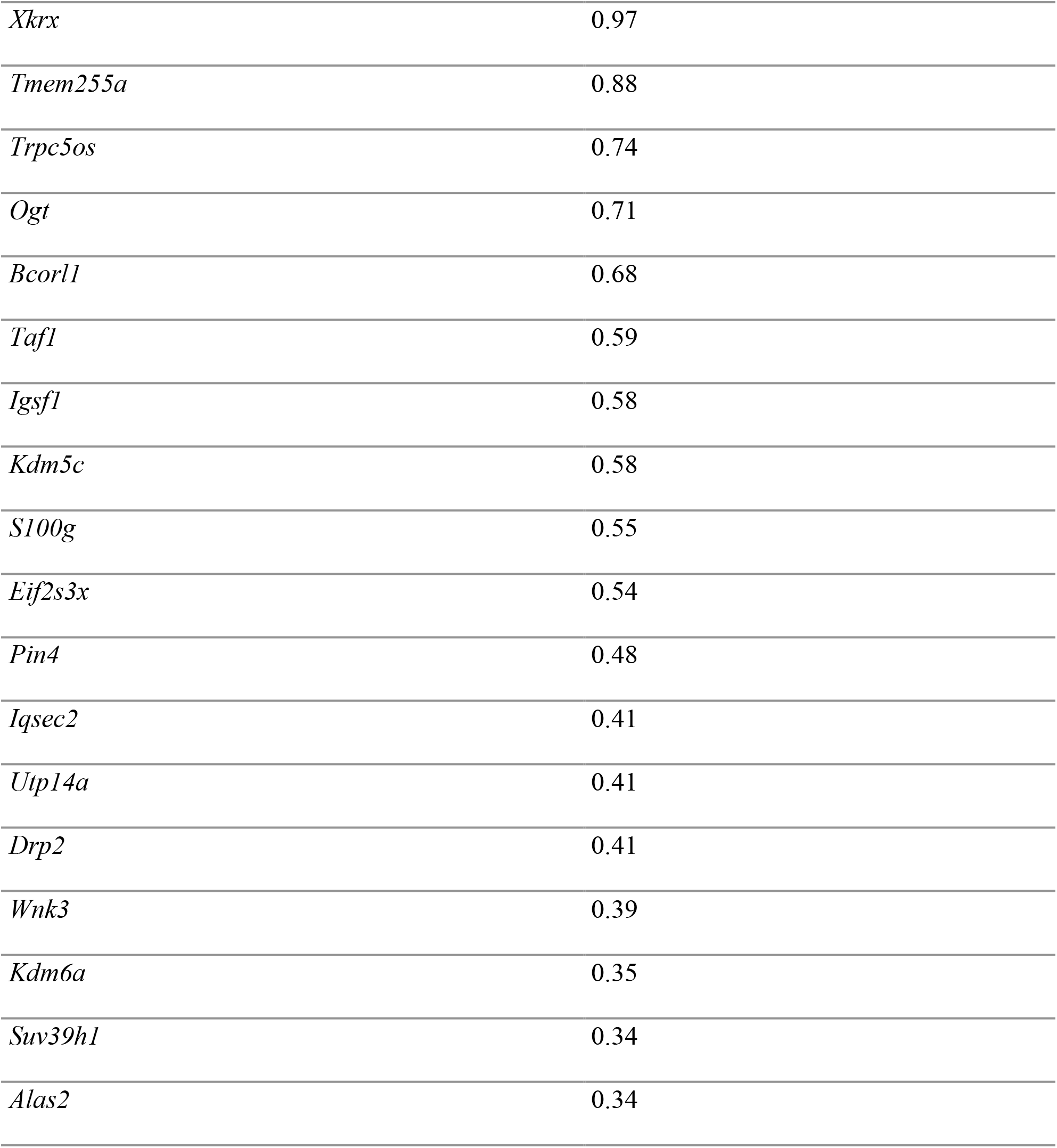
Top differentially expressed genes on the X chromosomes (*p* ˂ 0.05)

Gene ontology term enrichment analysis of autosomal DEGs with higher expression in male placentas revealed roles in regulating lactation (*Prl3d1*, *Prl2c2*), viral defense (*Apobec1*, *Ifi203*), and regulating responses to external stimuli (*C3ar1*, *Oprm1*; Figure 1E, left panel). In contrast, autosomal DEGs with higher expression in female placentas were enriched for roles in SLC-mediated transmembrane transport (*Slc22a1*, *Slc39a8*), tube morphogenesis (*Ackr3*, *Fgf18*), and gland development (*Cebpa*, *Ephb3*; Figure 1E, middle panel). X chromosome DEGs were notably enriched in genes encoding chromatin modifying enzymes (*Kdm5c*, *Xist*) and regulating small GTPase-mediated signal transduction (*Amot*, *Ogt*), actin cytoskeleton organization (*Arhgap6*, *Shroom2*), non-coding RNA processing (*Suv39h1*, *Ftsj1*), and cellular macromolecule biosynthesis (*Alas2*, *Eif2s3x*; Figure 1E, right panel).

These results reveal significant transcriptomic differences between male and female late-gestation mouse placentas. Notably, these differences extend beyond genes located on the sex chromosomes, indicating comprehensive impacts on gene expression throughout the autosomal genome.

### Late-gestation mouse placentas display DNA methylation sex differences

To further investigate sex-specific molecular differences in late-gestation mouse placentas, we generated genome-wide DNA methylation profiles of the same six E18.5 placenta samples using Methyl-Seq. After applying thresholds (*e.g*., minimum two CpGs, 100-bp tiles, 10× coverage in all samples), we identified 756638 tiles covering 2.4 million CpGs (Figure S3). Although the global DNA methylation patterns on these tiles were generally similar between male and female placentas (Figure S3A), we observed higher mean methylation values in male placentas (38.3%) than in female placentas (37.6%; Figure S3B). This trend was maintained across all autosomes (38.4% vs. 37.7%); however, the X chromosomes exhibited higher mean methylation levels in female placentas (29.9%) than in male placentas (29.0%; Figure S3C). These differences caused notable shifts in the percentages of tiles in different methylation categories (*e.g.,* 0–10% to 90–100%) between both the autosomal and X chromosomes of male and female placentas (Figure S3D–E).

We next analyzed the tiles to identify differentially methylated regions (DMRs; 100-bp tiles, > 10% increase or decrease, *q* < 0.01) in the genomes of male and female placentas. On autosomal chromosomes, we found 1705 DMRs (0.2% of the 745699 tiles) between male and female placentas (Figure 2A–C; Table S1). Of these, 73% (*n* = 1251) and 27% (n = 454) exhibited higher methylation levels in male and female placentas, respectively. Considerably more DMRs (*n* = 3756; 34% of 10939 tiles) were observed on the X chromosomes, with 66% (*n* = 2491) and 34% (*n* = 1265) displaying higher methylation levels in female and male placentas, respectively (Figure 2A–C; Table S1). Most autosomal (86%) and X chromosomal (72%) DMRs had 10–20% changes in their methylation levels between male and female placentas (Figure 2D). Only 2% of the autosomal DMRs and 4.5% of the X chromosomal DMRs had a > 30% change in methylation between the sexes.

**Figure 2.**
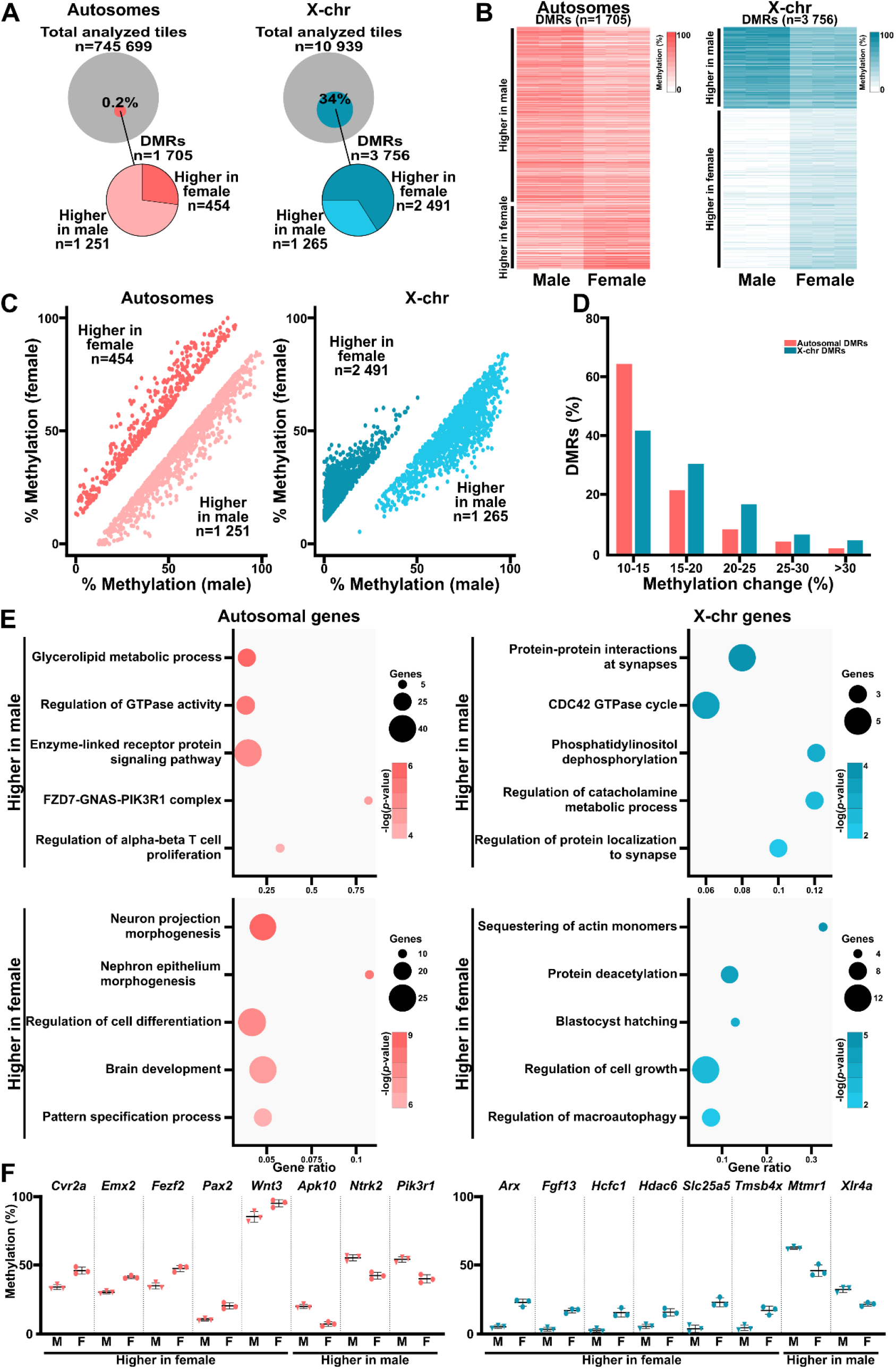
DMRs are present throughout the genomes of late-gestation mouse placentas. A) Schematic of the analyzed tiles in autosomes (left) and X chromosomes (right). DMRs exhibiting higher methylation levels in male or female placentas are indicated. **B)** DNA methylation levels of autosomal (left) and X chromosome (right) DMRs in male and female E18.5 placentas. Samples are clustered by methylation level. **C)** DMRs located on autosomes (left) and X chromosomes (right). Dark colors represent regions with ≥ 10% increased methylation in female placentas (higher in female), while light colors represent regions with ≥ 10% decreased methylation in female placentas (higher in male). **D)** The proportions of autosomal and X chromosomal DMRs that caused DNA methylation level changes of various magnitudes. **E)** The top five pathways enriched in autosomal (left) and X chromosomal (right) genes with DMRs that result in higher methylation levels in male (top) and female (bottom) placentas. **F)** DNA methylation levels of autosomal (left) and X chromosomal (right) DMRs associated with the top five enriched pathways in male and female placentas.

X chromosome DMRs with higher methylation in female placentas had average methylation levels below 50% in both sexes, while DMRs with higher methylation in male placentas predominantly displayed > 50% methylation (Figure 2C). This distribution was not observed for autosomal DMRs (Figure 2C). Some example DMRs are shown in Figure S4A, including *Cflr2*, which is involved in cell proliferation and hematopoietic system development; *Bcl2l11*, which is implicated in apoptosis, *Morf4l2*, which is predicted to play a role in heterochromatin assembly, and *Xist*, a well-known X chromosome inactivation factor. The top 20 DMRs on autosomes and X chromosomes are listed in Tables 3 and 4, respectively. Most DMRs were located in intronic and intergenic regions (76% and 56% on the autosomes and X chromosomes, respectively; Figure S4B). However, 26% of DMRs on the X chromosomes were located in promoter regions. We also observed methylation level changes across various genomic features on both the autosomes and X chromosomes (*e.g*., introns, promoters, transcriptional start sites (TTSs), 3′ and 5′ untranslated regions (UTRs), and intergenic regions; Figure S4C).

**Table 3.** Top autosomal DMRs.

| Gene symbol | Chromosome | Annotation | Change in methylation |
| --- | --- | --- | --- |
| Increased in male placentas |  |  |  |
| *Ube2k* | 5 | Promoter-TSS | -36.70 |
| *Tiam2* | 17 | Intron | -35.97 |
| *Map1b* | 13 | Intron | -35.44 |
| *Ccdc116* | 16 | 3' UTR | -35.31 |
| *Pygb* | 2 | Intron | -35.00 |
| *Ccdc60* | 5 | Intron | -34.52 |
| *Snord116l2* | 7 | Intron | -33.61 |
| *Lrpprc* | 17 | Intron | -33.26 |
| *Jdp2* | 12 | TTS | -32.69 |
| *Col6a3* | 1 | Intron | -32.53 |
| *Slc36a3* | 11 | Exon | -31.99 |
| *Ddr1* | 17 | Intron | -30.82 |
| *Mrpl45* | 11 | 3' UTR | -29.56 |
| *Alg6* | 4 | Intron | -29.31 |
| *Mir218-2* | 11 | Intron | -28.90 |
| *Trhr2* | 8 | Intron | -28.86 |
| *Tssc4* | 7 | Exon | -28.82 |
| *Krtdap* | 7 | TTS | -27.75 |
| *Gip* | 11 | Exon | -27.73 |
| *Nuak1* | 10 | Intron | -27.59 |
| **Increased in female placentas** |  |  |  |
| *Proz* | 8 | TTS | 38.15 |
| *Tmem267* | 13 | Promoter-TSS | 36.91 |
| *Cdk5rap1* | 2 | Intron | 34.71 |
| *Stam2* | 2 | Intron | 34.66 |
| *Cdk5rap1* | 2 | Exon | 33.96 |
| *Clip4* | 17 | Intron | 33.56 |
| *Cacng3* | 7 | Promoter-TSS | 32.84 |
| *Tmem267* | 13 | Promoter-TSS | 32.32 |
| *Pbx1* | 1 | Intron | 32.25 |
| *Adamts2* | 11 | Intron | 30.96 |
| *Cdk5rap1* | 2 | Intron | 30.28 |
| *Dennd1c* | 17 | Promoter-TSS | 29.27 |
| *Tmem267* | 13 | Promoter-TSS | 29.02 |
| *Tmem267* | 13 | Promoter-TSS | 27.87 |
| *Mbp* | 18 | Intron | 27.65 |
| *Ppox* | 1 | TTS | 27.44 |
| *Phf19* | 2 | Intron | 25.47 |
| *Bcl2l11* | 2 | Intron | 25.26 |
| *Nckap5* | 1 | Intron | 25.05 |
| *Cdk5rap1* | 2 | Intron | 25.01 |

**Table 4.** Top X-chromosome DMRs.

| Gene symbol | Annotation | Change in methylation |
| --- | --- | --- |
| **Increased in male placentas** |  |  |
| *Firre* | Intron | -47.74 |
| *Firre* | Intron | -46.93 |
| *Firre* | Intron | -46.80 |
| *Xist* | Non-coding | -45.23 |
| *Firre* | Intron | -43.31 |
| *Firre* | Intron | -42.18 |
| *Xist* | Non-coding | -40.44 |
| *Firre* | Intron | -40.22 |
| *Firre* | Intron | -39.42 |
| *Xist* | Non-coding | -39.07 |
| *Xist* | Non-coding | -38.95 |
| *Ikbkg* | Intron | -38.63 |
| *Firre* | Intron | -38.53 |
| *Mir3620* | Intron | -38.31 |
| *Xist* | Promoter-TSS | -37.99 |
| *Firre* | Intron | -37.88 |
| *Firre* | Intron | -37.55 |
| *Firre* | Intron | -36.76 |
| *Firre* | Intron | -36.44 |
| *Firre* | Intron | -36.39 |
| **Increased in female placentas** |  |  |
| *Morf4l2* | Intron | 39.94 |
| *Pja1* | Intron | 39.38 |
| *Msl3* | Intron | 36.36 |
| *Efnb1* | 5' UTR | 36.02 |
| *Taf7l* | Intron | 35.63 |
| *Amer1* | Intron | 35.58 |
| *Ddx3x* | Intron | 35.23 |
| *Pja1* | Intron | 34.69 |
| *Atp6ap1* | Intron | 33.83 |
| *Las1l* | Intron | 32.74 |
| *Pgk1* | Promoter-TSS | 32.59 |
| *Sox3* | 3' UTR | 32.54 |
| *Mir1970* | 5' UTR | 31.52 |
| *Cnksr2* | Intron | 31.30 |
| *Sox3* | Exon | 31.29 |
| *Rab33a* | Promoter-TSS | 30.96 |
| *Msl3* | Intron | 30.44 |
| *Efnb1* | Exon | 30.43 |
| *Rai2* | Intron | 30.36 |
| *Gk* | Promoter-TSS | 30.31 |

To examine the potential biological functions of these DNA methylation differences, we performed gene ontology term enrichment analyses on all DMRs located in gene-associated features like promoters, introns, and exons. For autosomal DMRs with higher methylation levels in male placentas (*n* = 735; in 672 unique genes such as *Apk10, Ntrk2,* and *Pik3r1*), the top pathways were related to metabolic processes and signaling pathways (Figure 2E). Conversely, DMRs with higher levels in female placentas (*n*= 303; in 254 unique genes such as *Cvr2a*, *Emx2,* and *Pax2*), were linked to brain development, cell differentiation/specification, and morphogenesis (Figure 2E). X chromosome DMRs with higher methylation levels in male placentas (*n* = 693; in 232 unique genes such as *Mtmr1* and *Xlr4a*) were associated with metabolic processes, protein interactions, and synaptic localization (Figure 2E). X chromosome DMRs with higher methylation in female placentas (*n* = 2237; in 430 unique genes such as *Arx, Fgf13,* and *Hdac6)* were associated with protein deacetylation and regulating cell growth and macroautophagy. The DNA methylation levels of selected gene-associated DMRs are shown in Figure 2F.

Together, these analyses reveal significant sex differences in DNA methylation profiles across the autosomes and X chromosomes of male and female E18.5 mouse placentas, despite their similar overall methylation status. These changes occur in critical genes and could impact their regulation during mouse placenta development. This underscores the importance of considering sex variations when studying placental methylation patterns.

### DMRs are enriched in CpG islands on the X chromosome but are located away from them on autosomes

To investigate the genomic contexts of DMRs in E18.5 placentas, we examined the annotations and CpG densities of autosomal and X chromosome DMRs. X chromosome DMRs had higher CpG enrichment per tile than autosome DMRs (4.8 vs. 3.4 CpGs per tile, *p* < 0.0001; Figure 3A). Of the autosomal DMRs, 3% (49 DMRs) contained > 10 CpGs per tile (maximum: 19 CpGs). X chromosome DMRs also contained up to 19 CpGs per tile; however, 11% (403 DMRs) contained > 10 CpGs. A significantly higher percentage of DMRs were present in high-density CpG islands (35%) and their immediate flanking regions (CpG shores; 33%) on X chromosomes than on autosomes (islands: 6%; shores 18%; *p* ˂ 2.2e^-16^; Figure 3B). Autosomal DMRs were enriched in regions away from CpG islands (open sea; autosomes: 69%; X chromosome: 29%; *p* ˂ 2.2e^-16^). The average DNA methylation levels in CpG island DMRs were higher in female placentas than in male placentas for both autosomal and X chromosomal DMRs (Figure 3C). However, in the CpG shelves, the average DMR methylation levels were higher in male placentas, averaging close to 50% methylation on both autosomes and X chromosomes.

**Figure 3.**
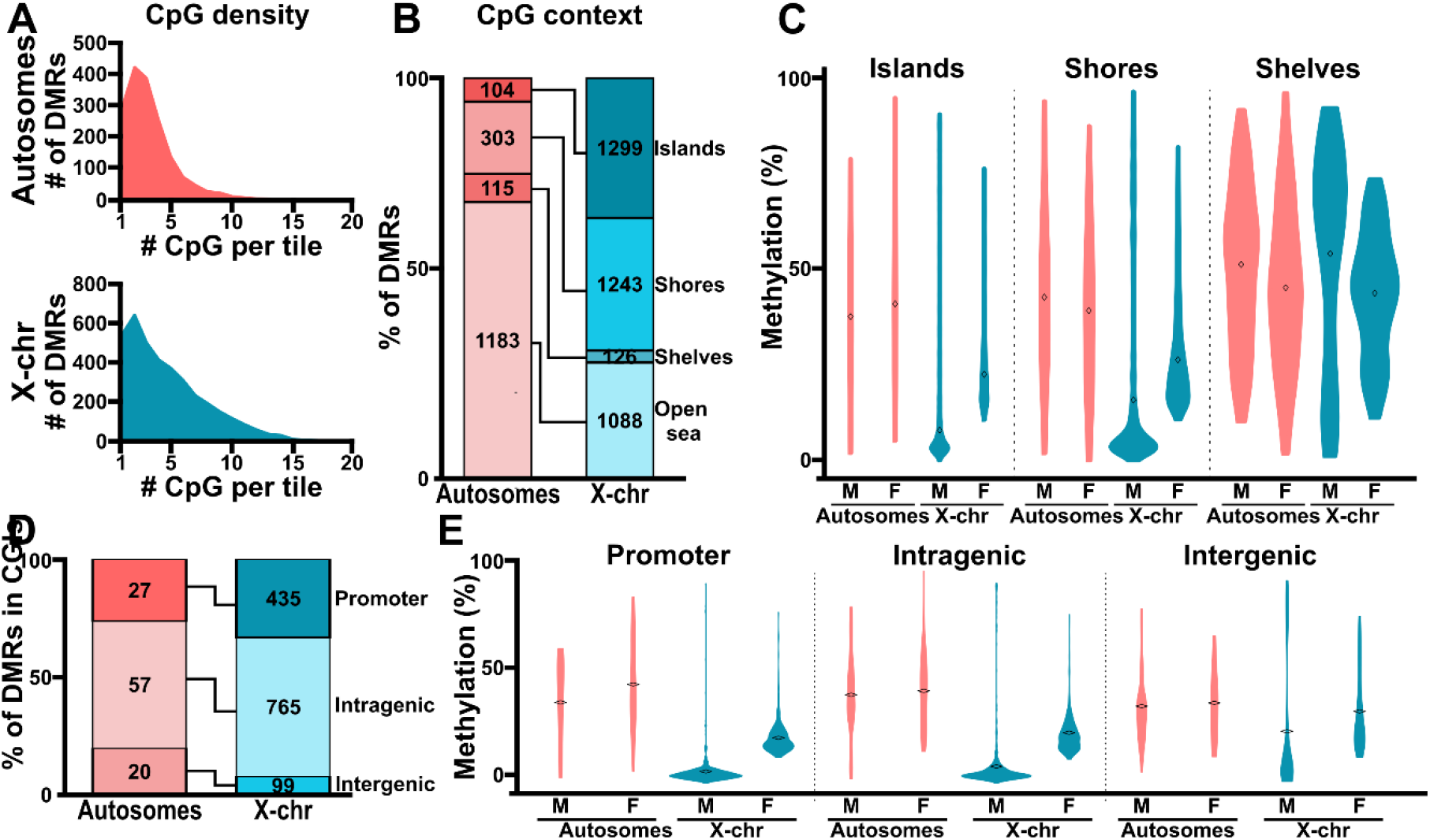
DMRs have divergent distributions on the autosomes and X chromosomes. A) The frequency distributions of autosomal (top) and X chromosome (bottom) DMRs at various CpG densities (*i.e*., numbers of CpGs per analyzed tile) in male and female E18.5 placentas. **B)** DMR distributions in male and female placentas based on their proximity to CpG islands (CGIs). Proximal regions are defined as shores (up to ±2 kb from a CGI), shelves (± 2–4 kb from a CGI), and open seas (± 4 kb or more from a CGI). **C)** DNA methylation levels on autosomal and X chromosome DMRs in male and female placentas by CGI proximity. **D)** Distributions of CGI-based DMRs into various genomic regions in male and female placentas. **E)** DNA methylation levels of CGI-based autosomal and X chromosome DMRs located in promoter, intragenic, and intergenic regions in male and female placentas.

Since CpG islands can function as TSSs and play key roles in regulating gene expression, we annotated the CpG island-associated DMRs to genomic features. We observed that 26% (*n* = 27) of autosomal DMRs and 33% (*n* = 435) of X chromosomal DMRs were in CpG islands within promoter regions (Figure 3D). The majority of CpG island DMRs (autosomes: 54%, *n* = 57; X chromosomes: 59%, *n* = 765) were located within genes (*i.e*., in intragenic regions). For CpG island DMRs present in promoters, intragenic, and intergenic regions, DNA methylation levels were higher in female placentas compared to male placentas (Figure 3E), with the average methylation level being lower on the X chromosomes.

Overall, DMRs were mainly found in CpG-poor regions on autosomes but in CpG-rich regions on the X chromosomes. However, the normally observed inverse correlation between DNA methylation and CpG density was not particularly obvious for DMRs located in autosomal CpG-rich promoters compared to their counterparts on the X chromosomes, suggesting that the higher methylation levels play a regulatory role during development.

### Associations between sex-specific DNA methylation and gene expression patterns in E18.5 mouse placentas

Our findings provide compelling evidence of sex-specific DNA methylation and gene expression levels in E18.5 placentas. To assess the associations between these variations, we compared the datasets. First, we annotated DMRs located in promoters or intragenic regions to unique genes. On autosomes, we identified 58 DMRs associated with 34 unique genes, while on the X chromosomes, 111 DMRs were linked to 18 genes. Next, we overlapped these DMR-associated genes with the identified DEGs (Figure 4A). This analysis identified 52 genes with 169 associated DMRs that displayed sex differences in both DNA methylation and gene expression profiles. Of these, 34 (linked to 58 DMRs) were autosomal, while 18 (linked to 111 DMRs) were located on the X chromosomes.

**Figure 4.**
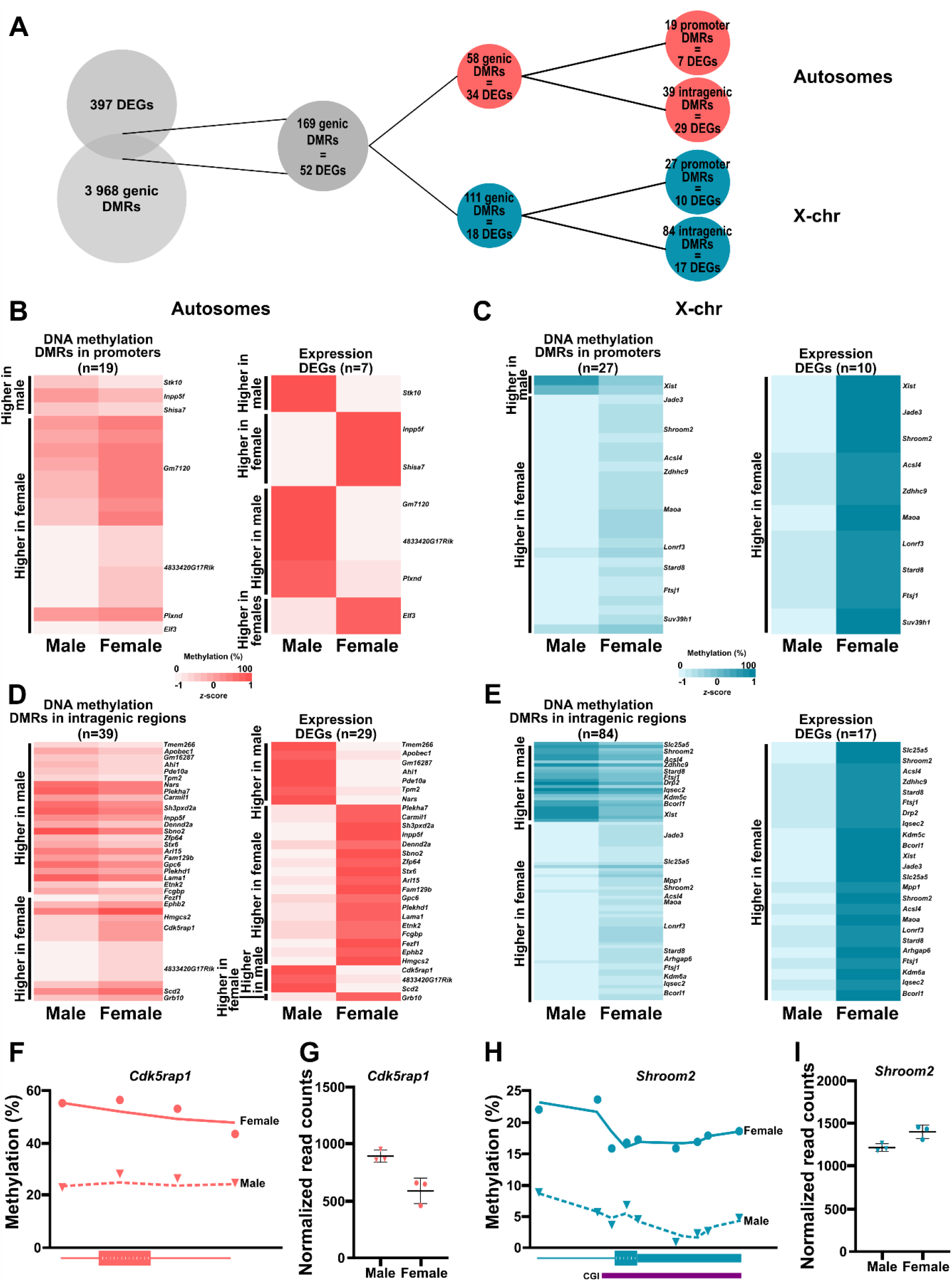
Correlations between gene expression and DNA methylation changes in the E18.5 mouse placenta. A) Overlap between DEGs and DMRs on the autosomes and X chromosomes and their locations in promoters and intragenic regions. **B)** and **C)** DNA methylation (left) and gene expression (*z*-scores; right) levels of promoter and gene regions with significant DNA methylation and expression sex differences on the autosomes (**B**) and X chromosomes (**C**) of E18.5 placentas. **D)** and **E)** DNA methylation (left) and gene expression (*z*-scores; right) levels of intragenic regions (excluding promoters) with significant DNA methylation and expression sex differences on the autosomes (**D**) and X chromosomes (**E**) of E18.5 placentas. **F)** *Cdk5rap1* DNA methylation levels in male and female placentas. Dots represent the mean methylation levels of individual DMRs in each sex. **G)** Normalized read counts for *Cdk5rap1* in male and female placentas. **H)** *Shroom2* DNA methylation levels in male and female placentas. Dots represent the mean methylation levels of individual DMRs in each sex. **I)** Normalized read counts for *Shroom2* in male and female placentas.

We also examined the distributions of these DMRs, specifically whether they were present in promoters or other intragenic regions (Figure 4B). On autosomes, we observed direct correlations between promoter DMRs and DEGs. For instance, genes like *Inpp5f*, *Shisa7, Gm7120, 4833420G17Rik*, and *Plxnd* displayed higher methylation levels in one sex and higher expression in the other (Figure 4B). However, this correlation was not observed for *Stk10* and *Elf3* (Figure 4B). On the X chromosomes, we observed this association only for *Xist*, which had higher methylation in male placentas and higher expression in female placentas. The remaining nine X chromosome promoter DMRs had higher methylation and gene expression levels in female placentas (Figure 4B). While the negative correlation between promoter DNA methylation and transcriptional expression is well documented (43, 44), the role of intragenic methylation is less clear. However, previous studies have shown positive associations between DNA methylation within intragenic regions and gene expression in various contexts (45–47). In our datasets, we identified 46 DEGs (29 on autosomes, 17 on X chromosomes) also displaying intragenic DMRs (Figure 4D–E). On autosomes, intragenic DMRs with higher DNA methylation levels in male or female placentas had both positive (male: *Tmem266*, *Apobec1*, *Pde10a;* female: *Ephb2*, *Hmgc2*, *Grb10*) and negative (male: *Plekha7*, *Carmll1*, *Zfp64*; female: *Cdk5rap1, 4833420G17Rik, Scd2*) correlations with gene expression. On the X chromosomes, intragenic DMRs with higher levels in either male or female placentas were associated with DEGs with increased expression in female placentas. Example associations between promoter and intragenic DMRs and gene expression are shown in Figures 4F–4I.

### DNA methylation and expression sex differences in genes essential for placental development

To assess the potential significance of DMRs in placental development and function, we conducted an overlap analysis between all DMRs and a curated list of 205 published essential placental genes (Table S2). Of these, 192 genes had sufficient sequencing coverage for downstream analyses, representing 7360 tiles in promoter and intragenic regions after excluding intergenic regions (Figure S5A). There were 47 DMRs (12 autosomal, 35 X chromosomal) associated with 16 unique essential placental development genes (10 autosomal, 6 X chromosomal; Figure S5B–D). Among the autosomal DMRs, 11/12 displayed higher methylation levels in male placentas, while 26/35 X chromosome DRMs were more methylated in female placentas (Figure S5B, E). Functional gene ontology term enrichment analysis of the 16 essential placental genes containing the 47 DMRs confirmed their pivotal roles in various processes related to placenta development and function (Figure S5F). Only 10/205 (∼ 0.5%) of the essential placenta genes showed significant expression sex differences (Figure S5G). *Serpine1* displayed higher expression in male placentas, while the rest (*Met*, *Pcld1*, *Ovol2*, *Havcr1*, *Etnk2*, *Gcm1*, *Dsg3*, and *Xist*) exhibited enhanced expression in female placentas. The higher expression of *Etnk2* (autosomal gene) and *Xist* (X chromosome gene) in female placentas also correlated with DMRs with higher methylation levels in male placentas.

Overall, our findings indicate that there are no substantial differences in the DNA methylation and gene expression patterns of crucial placental development genes between male and female placentas. This suggests that the underlying main control mechanisms driving placental development in late gestation are similar in both sexes. However, the presence of sex variations in DNA methylation or transcription patterns implies potential differences in the regulation of some placental functions during late pregnancy.

### Subsets of DMRs emerge during early placental development

To define the timing of the DMRs observed in late-gestation placentas, we compared the sex-based DMRs in mid-gestation (E10.5; *via* low-coverage reduced-representation bisulfite sequencing) and E18.5 placentas. The E10.5 placenta dataset contained 73 regions (100bp) that overlapped with the 1705 autosomal DMRs identified at E18.5 and 583 regions (100bp) that intersected with the 3756 E18.5 X chromosomal DMRs (Figure 5A–B). Of those overlapping regions, 19 autosomal regions were also differentially methylated (DMRs) when comparing male versus female placentas at E10.5. In E18.5 placentas, 6 autosomal DMRs exhibited higher methylation levels in male and 13 in female placentas, while at E10.5, methylation at these 19 DMRs was consistently higher in female placentas (Figure 5B). For most autosomal regions, we observed increased overall DNA methylation levels at E18.5 compared to E10.5 in all placentas, indicating a sex-independent developmental gain in DNA methylation (Figure 5B–C). One DMR, associated with *Vsx2*, displayed similar DNA methylation levels across all stages of development in both male and female placentas (Figure 5C). For other genes, (*Armc10*, *Grip1*, *Map7d1*), the DNA methylation levels at E10.5 and E18.5 closely matched only in female placentas. When we examined the 563 X chromosome DMRs, 348 (62%) were significantly different between male and female E10.5 placentas (Figure 5A, D). These regions all displayed higher DNA methylation levels in female placentas at both E10.5 and E18.5. Smoothed plots depicting multiple DMRs in *Arx*, *Bcor*, *Gpc3*, and *Hs6st2* are shown in Figure 5E.

**Figure 5.**
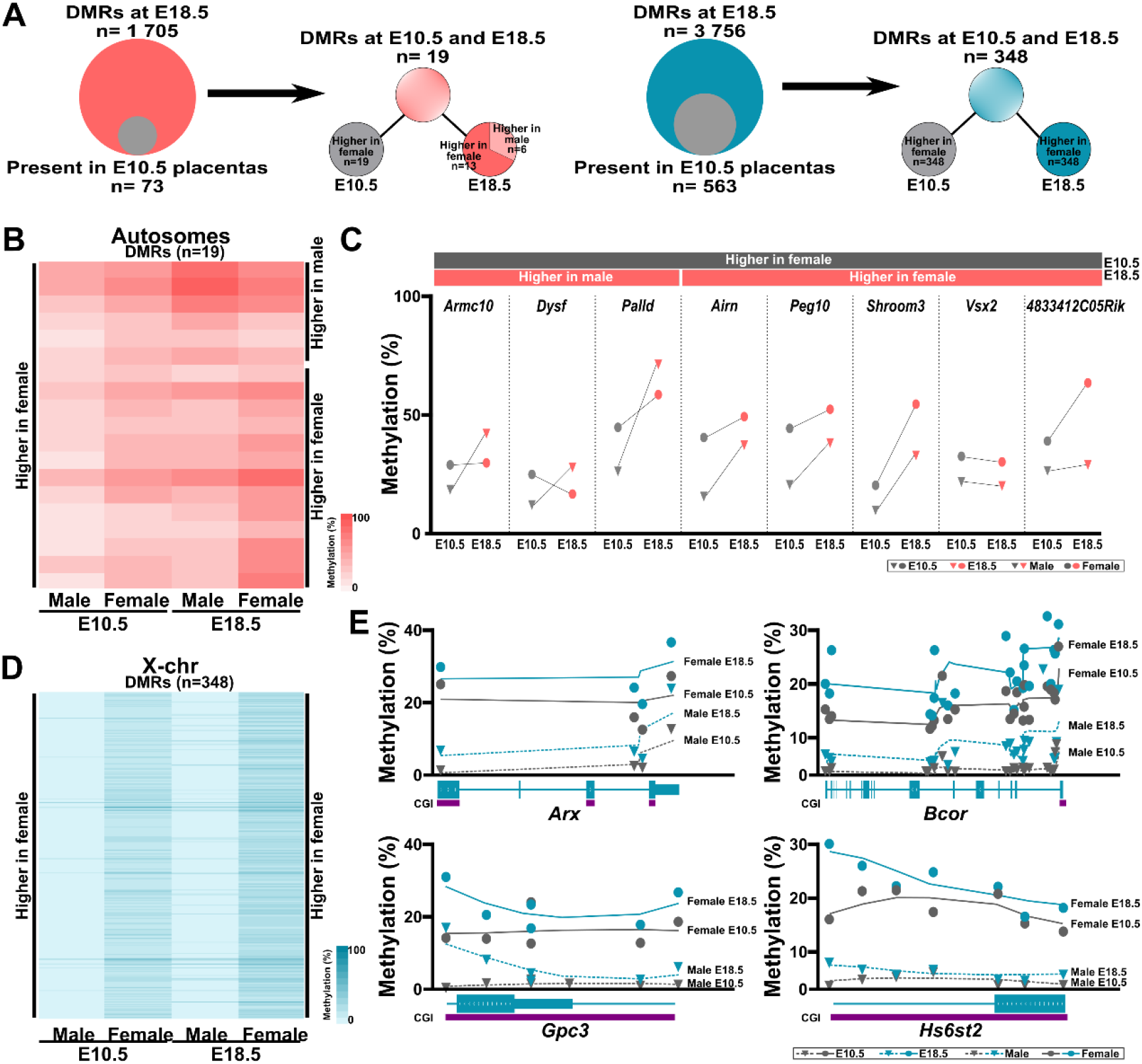
Selected DMRs between male and female E18.5 placentas are already established by mid-gestation. A) Schematic of the overlap between autosomal (left) and X chromosomal (right) DMRs at E10.5 and E18.5. The term ’Present’ designates 100bp tiles identified within the E10.5 placenta dataset with a coverage of at least 10X, exhibiting DMRs in E18.5 placentas **B)** DNA methylation levels of DMRs present on autosomal chromosomes at both E10.5 and E18.5 in male and female placentas. **C)** Mean DNA methylation levels of selected autosomal DMRs in male and female placentas at E10.5 and E18.5. **D)** DNA methylation levels of DMRs present on the X chromosome at both E10.5 and E18.5 in male and female placentas. **E)** DNA methylation profiles of X chromosome DMRs in the indicated genes.

These findings suggest that as the maturing mouse placenta (E10.5) develops and becomes more complex (e.g., cell types, organisation), DMRs arise. The presence of both consistent methylation levels across development and differential methylation patterns in specific genes highlights the intricate and selective nature of sex-specific epigenetic regulation in the placenta.

## DISCUSSION

In this study, our objective was to identify molecular sex differences in the gene expression and DNA methylation profiles of male and female late-gestation mouse placentas. Consistent with the intricacies of proper development and X chromosome inactivation, a significant proportion of the molecular changes occurred on the X chromosomes, which had particularly divergent DNA methylation profiles and displayed sex differences in gene expression. However, significant epigenetic and transcriptomic sex differences were also observed on the autosomes. These data demonstrate that sex differences are widespread in the mouse placenta genome, and that simply excluding sex chromosome-derived sequencing data is an inadequate means of mitigating their effects in -omics studies.

On autosomes, female placentas had more DEGs with increased expression, while male placentas had more DMRs with increased methylation levels. These differences were associated with various biological functions, consistent with the mounting evidence of a mechanistic link between placental pathways and growth sex differences and risks of pregnancy complications (6, 48, 49). Male placentas exhibited higher expression of genes associated with immune responses than female placentas, including DEGs involved in the responses to interferon α (*Ifit*, *Ifit2*), viruses (*Apobec1*, *Ifi203*), and other external stimuli (*C3ar1*, *Oprm1*). This suggests that male and female placentas could have differences in immune response. Notably, the male fetal-placental unit is more sensitive to maternal inflammation than the female unit (50). Studies have reported upregulated innate and adaptive immune responses in male placentas, including in cases of maternal SARS-CoV-2 infection (reviewed in (51)). Maternal infection during early pregnancy is a recognized prenatal risk factor for mental illness in the offspring, particularly in males (52, 53). A recent study examining the consequences of a high-fat diet during pregnancy, which creates a chronic inflammatory environment, highlighted the significant influence of sex in shaping distinct vulnerabilities and outcomes affecting the placenta, fetal brain, adult brain, and behavior (54). It remains uncertain whether increase expression of immune response genes in the male fetal-placental unit could confer potential advantages, such as protection against viral infections, or disadvantages, such as increased placental inflammation, higher risk of fetal growth restriction, or impaired placental function.

Genes with higher DNA methylation levels in male placentas were linked to metabolic processes and signaling pathways, while genes that were more highly methylated in female placentas were associated with brain development, cell differentiation/specification, and neuron morphogenesis. Sexual dimorphism contributes to slight variations in placental metabolism, which have been speculated to be adaptive responses triggered by the fetus to promote optimal growth and ensure healthy development. In a recent study using a mouse model (C57BL/6J mice, standard diet) similar to the one used here, Saoi *et al*. observed distinct metabolic differences between male and female E18.5 placentas. Specifically, the levels of intracellular metabolites related to fatty acid oxidation and purine degradation were higher in female placentas than in male placentas. This is consistent with our expression data showing that fatty acid transporter proteins like SLC27A1—which are critical for nutrient transport in the placenta—are more highly expressed in female placentas. Interestingly, we observed DMRs in neurodevelopmental genes, highlighting the connection between the placenta and the brain (the “placenta-brain axis”) (54, 55). Various studies have reported associations between brain development and DNA methylation sex differences in autosomal genes, with specific DNA methylation profiles related to both normal and complicated pregnancies. For instance, in a mouse model of neurodevelopmental disorder induced by exposure to polychlorinated biphenyls, Laufer *et al*. observed shared sex-specific DNA methylation alterations related to neurodevelopment and autism spectrum disorder that were shared by the fetal brain and placenta (56). In their study and in others, prenatal insults consistently impact male offspring more significantly than female offspring (6, 56, 57). Importantly, these studies provide evidence that placental DNA methylation profiles can predict brain DNA methylation profiles—and possibly the risk of neurodevelopmental disorders.

The sex differences in DNA methylation levels we observed within the placenta were consistent with those observed in humans, in that male placentas typically exhibited higher methylation levels at DMRs than female placentas (24, 58). Interestingly, it is the opposite of what occurs in most somatic tissues, where sex-associated DMRs tend to be more highly methylated in females (59–63). Furthermore, we observed that some sex differences in DNA methylation levels occurred as early as E10.5, suggesting that they emerge during early placental development. Notably, we also identified regions where methylation remains consistent within one sex across both time points, particularly in autosomal regions. This indicates that in certain regions, DNA methylation is stable in one sex throughout embryonic development but highly dynamic in the other. Our findings also align with sex differences observed in human placental DNA methylation profiles associated with gestational age (64). Further research will be required to fully comprehend the potential implications of these sex differences on fetal development in both healthy and complicated pregnancies.

Despite the thousands of DMRs observed on the X chromosomes, only a few dozen genes exhibited differential expression between male and female placentas. However, the increased expression of several X-linked chromatin-modifying enzymes and transcription factors (*e.g*., *Bcorl1*, *Jade3*, *Kdm5c*, *Kdm6a*, *Ogt*, *Suv39h1*, *Wnk3*, *Xist*, and *Zdhhc9*) in female placentas could profoundly influence their epigenetic landscape and contribute to the unique responses of male and female placentas to the maternal environment. While *Xist* is normally not expressed in male cells, other genes, such as *Ogt* (an O-linked N-acetylglucosamine (*O*-GlcNAc) transferase) and *Kdm5c* (a histone H3K4-specific demethylase) have been observed at higher baseline levels in female placentas due to their ability to escape X inactivation (65). In addition, OGT is selectively downregulated in male placentas from mothers with gestational diabetes (66), and mouse studies indicate that prenatal stress impacts OGT and O-GlcNAcylation levels more in males than in females (67). The finding that suboptimal environments can influence the expression of X-linked genes involved in key epigenetic regulatory processes reinforces the need to systematically acquire and analyze sex-specific measurements in transcriptomic and epigenetic placental studies, especially those investigating responses to adverse maternal environmental stimuli.

## PERSPECTIVES AND SIGNIFICANCE

There has been comprehensive research investigating how the placenta contributes to pregnancy complications and the development of future offspring, particularly focusing on sex-related differences. Yet, despite these efforts, our understanding of sex-based variations in the epigenetic (like DNA methylation) and transcriptomic characteristics of the late-gestation mouse placenta remains considerably limited. Our study uncovers significant sex differences in the DNA methylation and gene expression profiles of late-gestation mouse placentas, providing comprehensive lists of the affected regions and genes. Importantly, we observed changes not only on the X chromosome but also the autosomes, demonstrating the importance of accounting for sex differences and not assuming equivalency between males and females on non-sex chromosomes. These findings emphasize the importance of examining the impacts of sex-specificity in epigenetic and transcriptomic research, regardless of the species or developmental staged studied. Recognizing sex as a crucial biological variable will enhance our understanding of the intricate interplay between fetal sex, placental biology, and adverse maternal environmental stimuli, ultimately advancing our knowledge of reproductive health and improving pregnancy outcomes.

## Declarations

The authors have no conflicts of interest to declare.

## Ethics approval and consent to participate

All animal studies were approved by the CHU Ste-Justine Research Center *Comité Institutionnel de Bonnes Pratiques Animales en Recherche* under the guidance of the Canadian Council on Animal Care.

## Consent for publication

Not applicable

## Availability of data and materials

All datasets used will be publicly available via the Gene Expression Omnibus.

## Competing interests

The authors declare no competing interests.

## Funding

This work was supported by a research grant to SM from the Natural Sciences and Engineering Research Council of Canada. LML is supported by Canadian Institutes of Health Research scholarship. MBL is supported by a scholarship/fellowship from Fonds de Recherche du Québec - Santé (FRQS). ALA is supported by scholarships from Université de Montréal and Réseau Québécois en Reproduction. SM is supported by an FRQS-Junior 2 salary award.

## Authors’ contributions

LML and SM conceptualized the study. LML and MBL contributed to data acquisition. LML, ALA, AL, participated in data analysis. LML and SM wrote the manuscript. All authors read and approved the final manuscript.

## Acknowledgements

We thank the McGraw lab for critical comments and suggestions, the staff of the Centre de Recherche du CHU Sainte-Justine animal facility for their assistance, and High-Fidelity Science Communications for editing the manuscript.

---

**Supplemental Figure 1.**
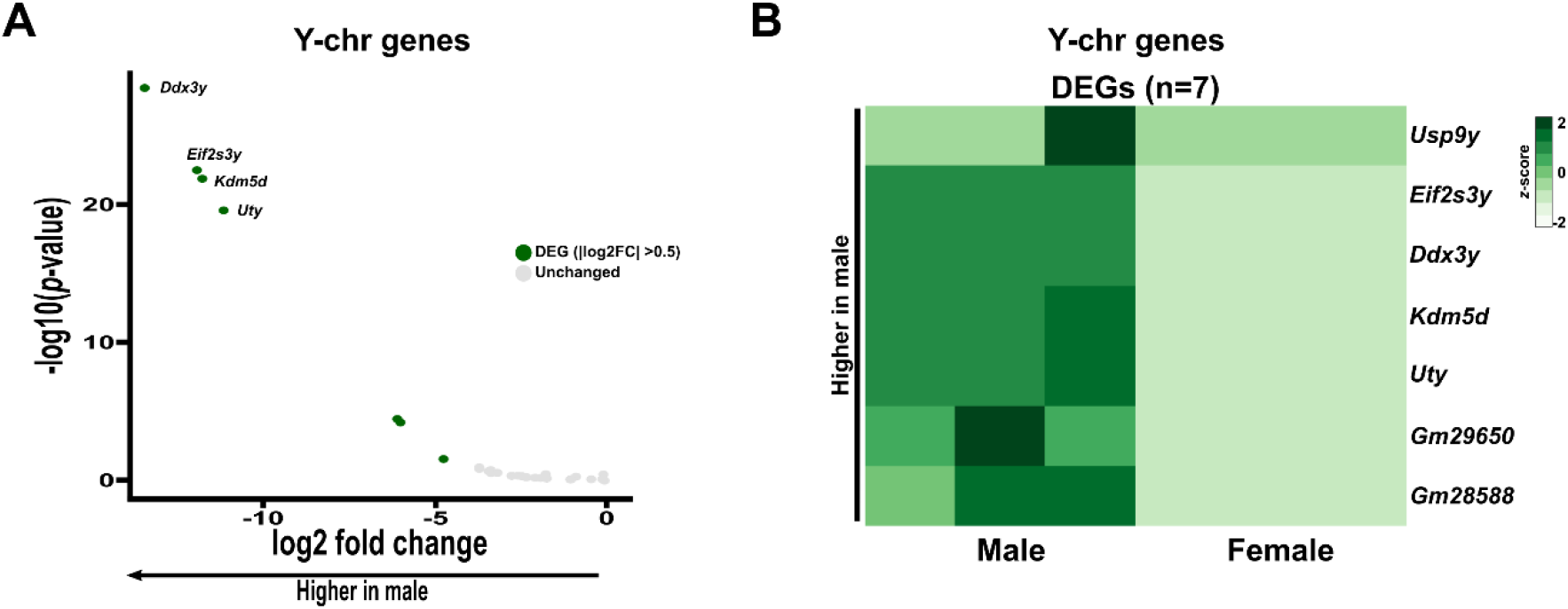
Expression of Y chromosome genes in male placentas. **A)** Differential expression analysis of Y chromosome genes in male and female E18.5 placentas (*n* = 121). Colored dots represent statistically significant DEGs (*p* < 0.05; *n* = 7) **B)** Expression levels (*z*-scores) of the Y chromosome DEGs.

**Supplemental Figure 2.**
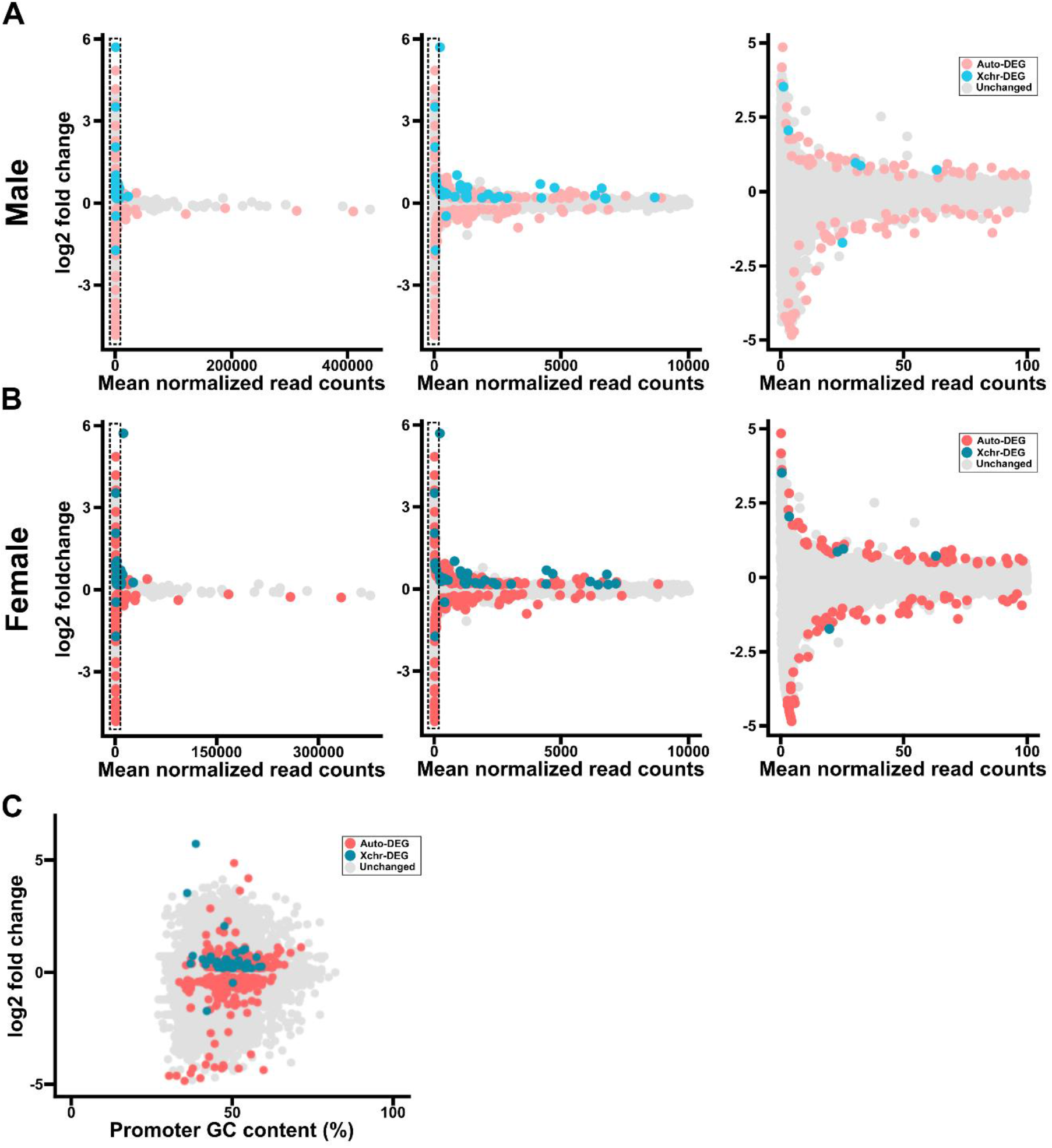
DEG abundance in male and female placentas. **A–B)** MA plot of the differential expression (log2 fold change) values of normalized read counts and their average expression levels in **A**) male and **B)** female E18.5 placentas. The plots include all analysed genes (left; *n* = 29480); genes with read counts > 10,000 (middle; *n* = 29165 and 29,160 in male and female placentas, respectively), and genes with read counts > 100 (right; *n* = 18160 and 18173 in male and female placentas, respectively). Colored dots represent significant DEGs on autosomes (pink) and X chromosomes (blue). The dotted black rectangle indicates the subset of the graph shown on the right. **C)** Differential expression values of each analyzed gene) in male and female placentas relative to their promoter’s GC content (%). Significant DEGs on autosomes and X chromosomes are shown in pink and blue, respectively.

**Supplemental Figure 3.**
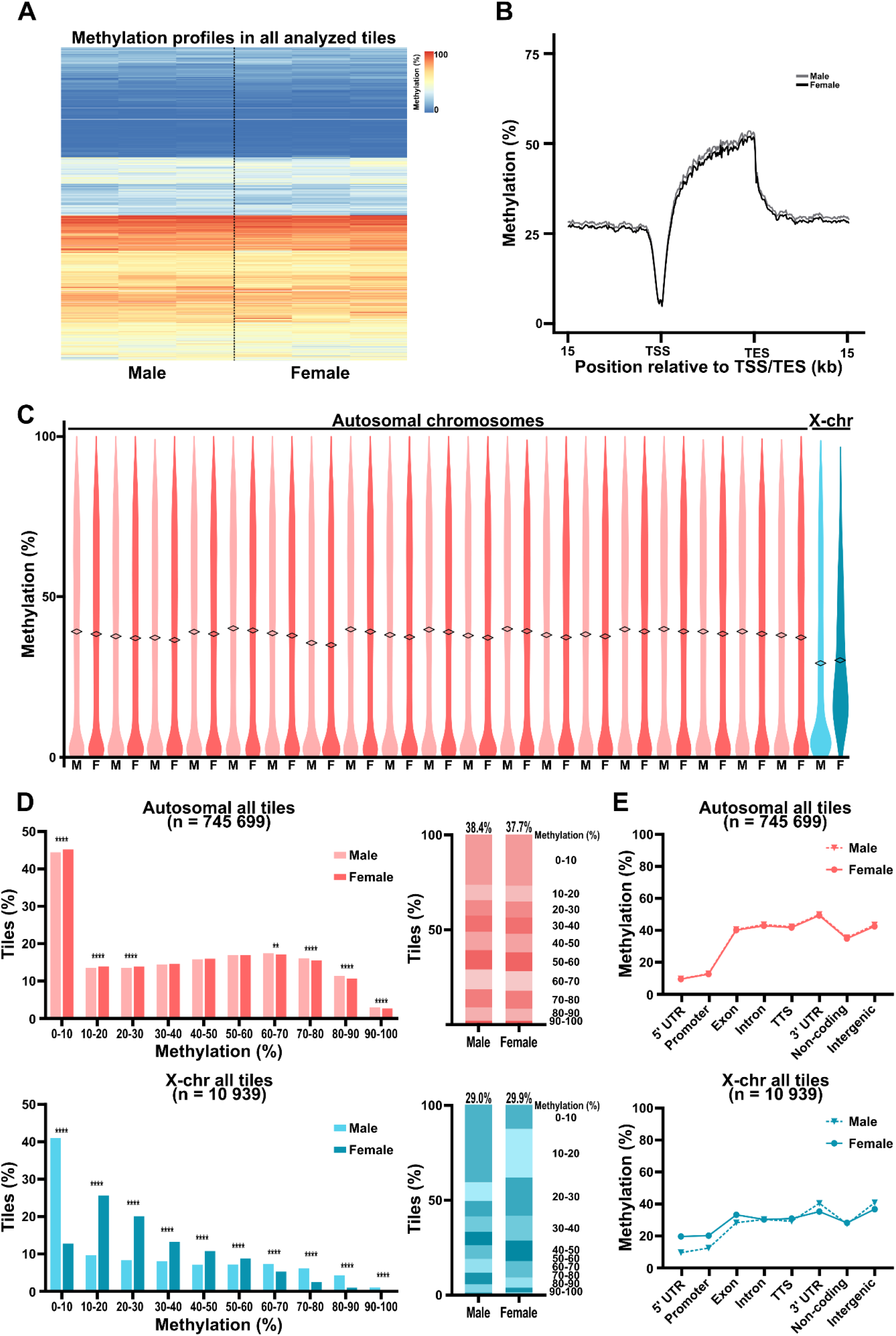
DMRs occur throughout the E18.5 placenta genome but are concentrated on the X chromosomes. **A)** DNA methylation levels of a random subset of tiles in the individual male and female placenta samples. **B)** Mean DNA methylation levels within ±15 kb of a transcriptional start site (TSS) or transcriptional end site (TES) in all analyzed tiles from male and female placentas. **C)** Distributions of the DNA methylation levels of various chromosomes in male and female placentas. Median DNA methylation values are indicated with diamonds. **D)** DNA methylation levels in tiles associated with autosomes (top) and X chromosomes (bottom) in male and female placentas. \*\*\*\**p* < 0.0001, \*\*\**p* < 0.001 by two-proportion*z*-test. **E)** Average DNA methylation levels in tiles associated with various genomic features on the autosomal (top) and X (bottom) chromosomes of male and female placentas.

**Supplemental Figure 4.**
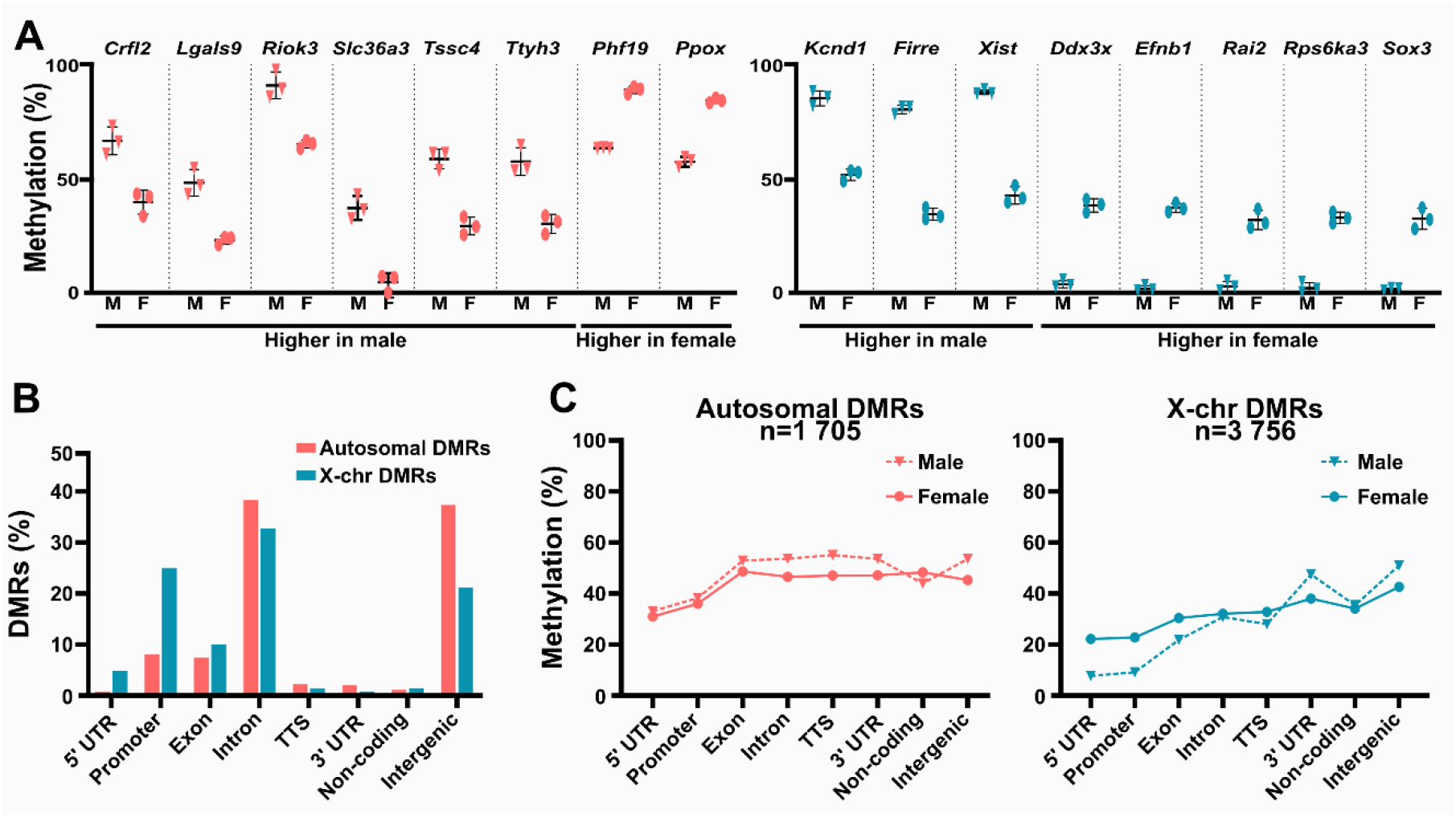
DNA methylation sex differences in various genomic elements. **A)** DNA methylation levels of top changed genes on the autosomes (left) and X chromosomes (right) of male and female placentas. **B)** Distributions of autosomal (pink) and X chromosome (blue) DMRs located in various genomic elements in male and female placentas. **C)** Average DNA methylation levels of DMRs in autosomes (left) and X chromosomes (right) based on their location’s genomic annotation.

**Supplemental Figure 5.**
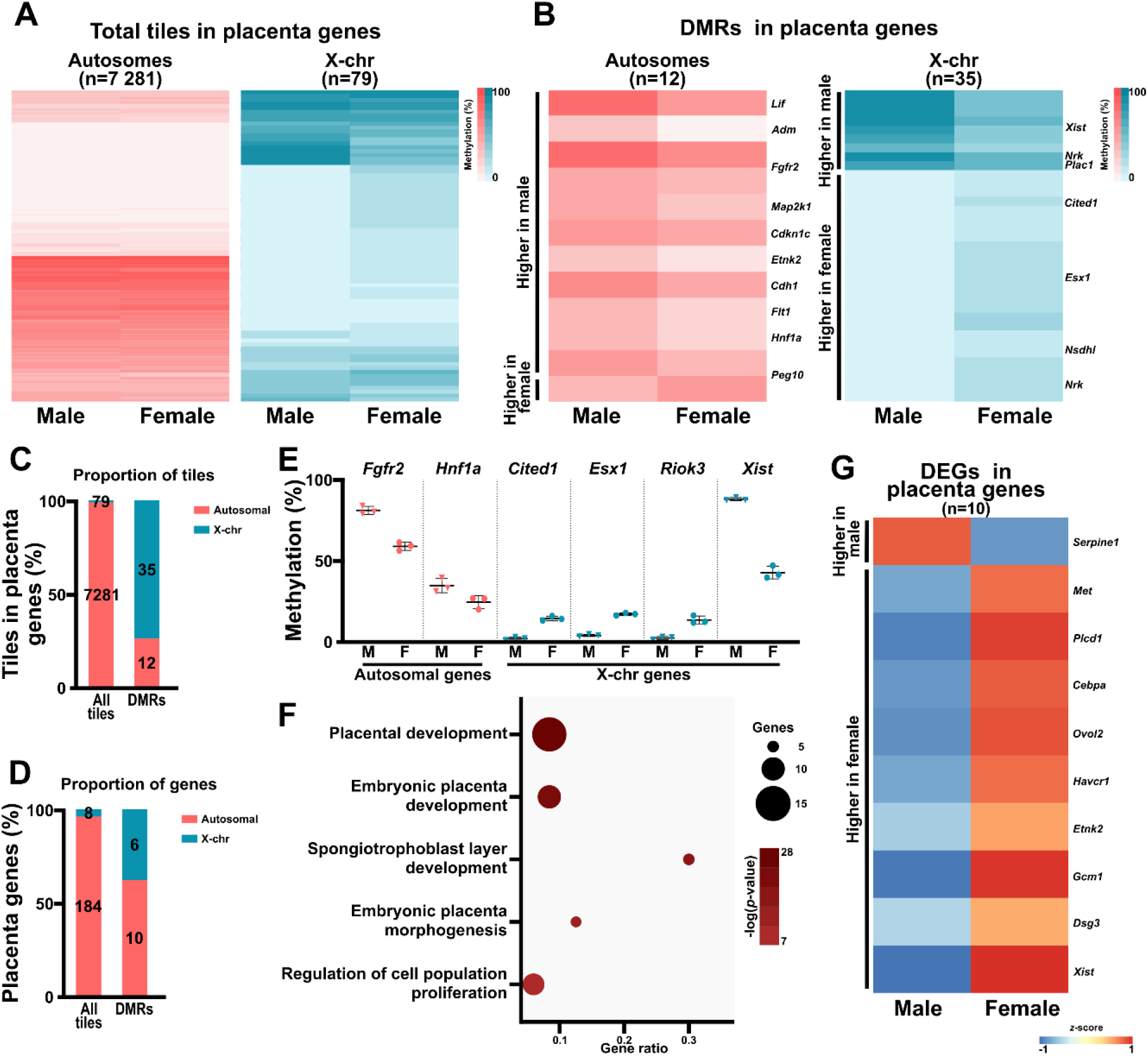
DNA methylation and expression patterns of genes essential for placental development. **A)** DNA methylation levels of autosomal (left) and X chromosomal (right) tiles associated with key placental developmental genes in male and female E18.5 placentas. **B)** DNA methylation levels of autosomal and X chromosomal DMRs associated with placental development genes in male and female placentas. **C)** Distributions of all analyzed tiles and DMRs located in essential placental development genes on the autosomes and X chromosomes. **D)** Distributions of individual placenta developmental genes on the autosomes and X chromosomes in the analyzed tiles and DMRs. **E)** DNA methylation levels of selected placenta developmental gene-related DMRs in male and female placentas. **F)** The top five pathways enriched for the placental development-associated DMRs in B). **G)** Expression levels (*z*-scores) of differentially expressed essential placental development genes.

## Notes

### Competing Interest Statement

The authors have declared no competing interest.

### Summary of Updates

Introduction was shorten. Modifications were made across the text to improve the message.

